# Genomic evidence of paternal genome elimination in the globular springtail *Allacma fusca*

**DOI:** 10.1101/2021.11.12.468426

**Authors:** Kamil S. Jaron, Christina N. Hodson, Jacintha Ellers, Stuart JE Baird, Laura Ross

## Abstract

Paternal genome elimination (PGE) - a type of reproduction in which males inherit but fail to pass on their father’s genome - evolved independently in six to eight arthropod clades. Thousands of species, including several important for agriculture, reproduce via this mode of reproduction. While PGE is well established in some of the clades, the evidence in globular springtails (Symphypleona) remains elusive, even though they represent the oldest and most species rich clade putatively reproducing via PGE. We sequenced genomic DNA from whole bodies of Allacma fusca males with high fractions (>27.5%) of sperm to conclusively confirm that all the sperm carry one parental haplotype only. Although it is suggestive that the single haplotype present in sperm is maternally inherited, definitive genetic proof of the parent of origin is still needed. The genomic approach we developed allows for detection of genotypic differences between germline and soma in all species with sufficiently high fraction of germline in their bodies. This opens new opportunities for scans of reproductive modes in small organisms.

## Introduction

The mechanism of reproduction varies considerably across the tree of life (Normark, 2003; Bachtrog et al., 2014). Historically, cytological comparisons of male and female karyotypes have been used to determine the mode of reproduction in a species. However, cytological studies are labour intensive and not all species have visible sex-specific karyotypes. As a consequence, many species still have undefined reproductive systems. On the other hand, genomic techniques have been successfully deployed to identify sex chromosomes in many taxa such as Diptera (Vicoso & Bachtrog, 2015; Anderson et al., 2020), and Lepidoptera (Fraïsse et al., 2017) and more recently to understand the exact form of parthenogenesis in species such as californian stick insects (Jaron et al., 2021), and bdelloid rotifers (Simion et al., 2021). Now, it is time to consider the ways we can use genomic techniques to study other modes of reproduction, such as paternal genome elimination.

Paternal genome elimination is a reproduction system in which males develop from fertilised eggs, but pass to the next generation only the maternally inherited haplotype (see Burt & Trivers, 2006 for an introduction to the topic). The inheritance pattern is exactly the same as in better known haplodiploidy (arrhenotoky), in which males develop from unfertilized haploid eggs, but mechanistically they represent very different reproductive systems. Similar to haplodiploidy, there are only a few known transitions to PGE (six to eight), and PGE clades are frequently very old and diverse. Thousands of arthropod species reproduce via some form of paternal genome elimination including human parasites (head and body lice), numerous agricultural pests (scale insects, Hessian fly, lucerne flea) and even pest control species (phytoseiid mites). However, the occurrence of PGE is likely significantly under-reported as it can be hard to confirm. It tends to occur in small arthropods that are poorly studied and hard to rear under laboratory conditions, making it challenging to study inheritance patterns. For example, PGE was only demonstrated in *Liposcelis* lice and human head and body lice (order Psocodea) very recently through genetic crosses tracking alleles over several generations (de la Filia et al., 2018; Hodson et al., 2017), even though meiosis was known to be unusual in lice for decades prior to this (Doncaster & Cannon, 1920; Cannon, 1922). Because of the difficulty of inheritance studies, many of the reported cases are based on indirect evidence, usually cytogenetic observations of unusual chromosome behaviour (**SM Table 1**).

Part of the reason PGE is difficult to identify, is that individual clades differ greatly in the mechanism of PGE, and hence require different types of evidence for confirmation (**Figure 1**). In all PGE species males develop from fertilised diploid eggs, and always exclusively transmit maternally inherited chromosomes to offspring. However, they differ in the processes leading to the elimination of paternal chromosomes. For example, in *Phytoseiidae* mites and some armoured scale insects, the paternal genome is completely eliminated early in embryogenesis in a process called embryonic PGE (Brown, 1965; Nelson-Rees et al., 1980). The fact that males are completely haploid soon after fertilisation makes this type of PGE easy to detect in genetic and cytological studies, although it can be hard to distinguish from true haplodiploidy. The two can be distinguished, however, via carefully designed phenotypic or irradiation crosses (Hoy, 1979; Helle et al., 1980), by cytology of early embryogenesis (Nelson-Rees et al., 1980), or by observing whether unfertilised eggs develop into males (Häußermann et al., 2020).

**Figure 1.**
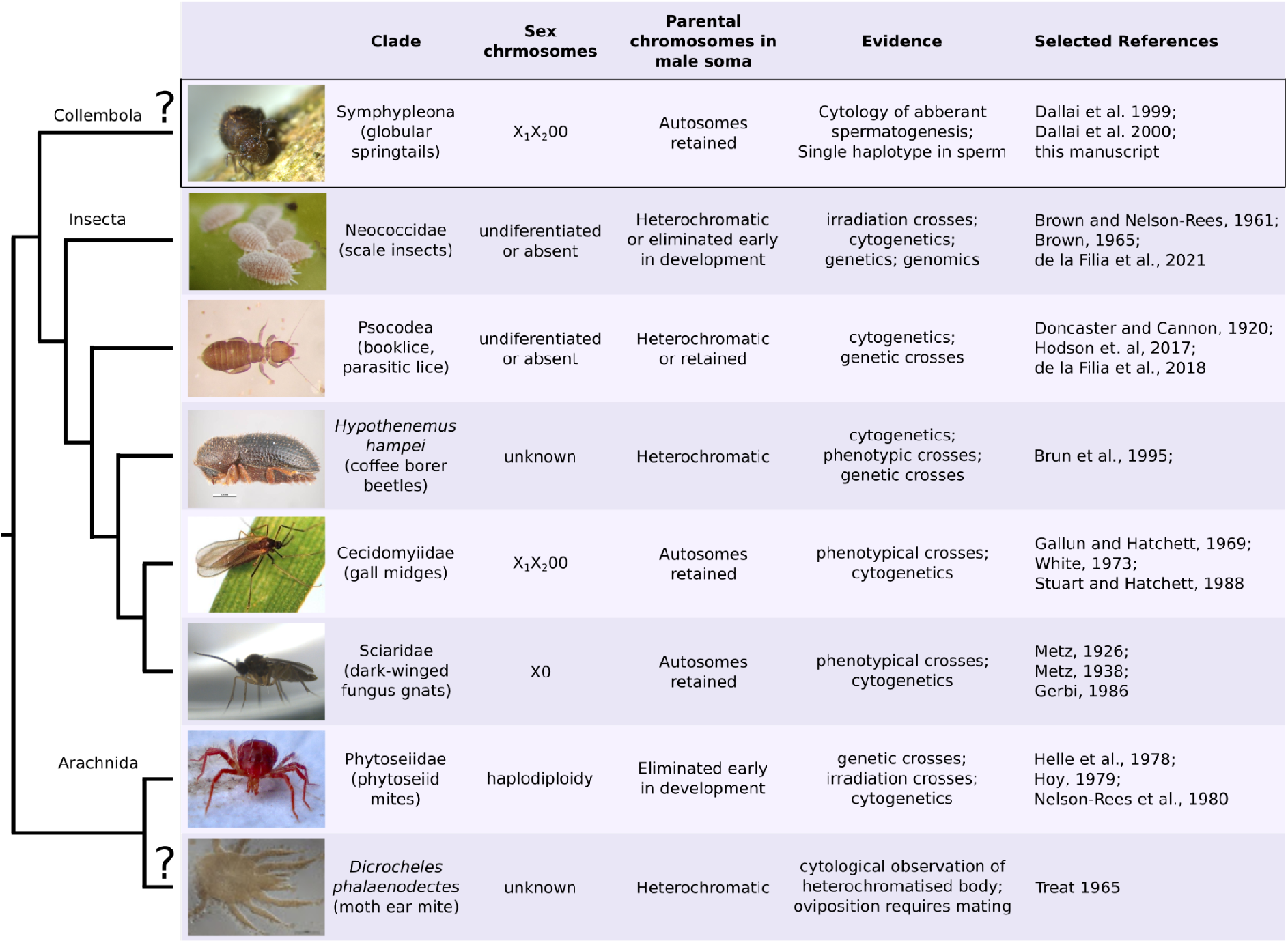
Clades with suggested paternal genome elimination (PGE) and evidence supporting it. The cladogram shows the phylogenetic relationships between putative (marked by “?”) and confirmed PGE clades. Although all PGE clades exhibit transmission dynamics where paternally inherited chromosomes are not transmitted to offspring through males, the sex chromosome system and the treatment/expression of paternally inherited chromosomes in male somatic cells can differ between and within clades. A more detailed list of relevant literature is in **SM Table 1.**. Image credits: mealybugs (scale insects) by Andrew J. Mongue, coffee borer beetles by Walker, K., phytoseiid mite by Mick Talbot.

In other types of PGE, males fully or partially retain their paternal genome throughout development and paternal chromosomes are excluded during spermatogenesis only, hence these types are known as germline PGE. While paternal chromosomes are retained, they form a dense heterochromatic bodies at the periphery of the cell nuclei for most scale insects (Brown, 1965; Ross et al., 2012), the coffee borer beetle (Brun et al., 1995), booklice (Hodson et al., 2017), and potentially in some Leapideae mites (Treat, 1965). This distinctive feature is not a formal test of PGE, but allows potential PGE species to be easily detected using cytological observation. It also means that males in these clades are mostly haploid in terms of gene expression, despite their diploid karyotype (Brun et al., 1995; de la Filia et al., 2021). A combination of embryonic and spermatogenic elimination is found in two dipteran families: fungus gnats, and gall midges. Males of these clades exclude one or two paternal chromosomes in early embryogenesis (usually referred to as X chromosomes), while retaining all other chromosomes in their soma. The remaining paternal chromosomes are lost during meiosis with a monopolar spindle which excludes paternal chromosomes from sperm. In fungus gnats and gall midges, it has been shown by crosses that all the eliminated chromosomes in both embryogenesis and spermatogenesis are of paternal origin (Metz, 1926, 1938; Gallun & Hatchett, 1969; Stuart & Hatchett, 1988). Finally a similar type of PGE has been suggested to occur in globular springtails. However the evidence is solely based on unusual chromosome behaviour and no inheritance studies are available.

Globular springtails are a large and species-rich order with enormous importance for soil ecology (Hopkin, 1997). Their karyotype consists of four to five autosomes and two sex chromosomes referred to as X_1_ and X_2_ (Dallai et al., 2000, 2004). Male globular springtail zygotes are initially fully diploid, but during very early embryogenesis they eliminate one copy of the X_1_ and X_2_ chromosomes (Dallai et al., 2000). Then during meiosis I of spermatogenesis the two X chromosomes co-segregate (i.e. are transmitted together), hence half of the secondary spermatocytes carry all six chromosomes and the other half contain the four autosomes only (**Figure 2**). The X chromosome-lacking spermatocytes immediately degenerate, and only the spermatocytes with the complete chromosome set undergo a second meiotic division to form two haploid spermatids (Dallai et al., 2000). In contrast to spermatogenesis of the majority of described species, only two of the potential four meiotic products yield in functional sperm, which is the reason this process is referred to as aberrant spermatogenesis. In a series of papers, Dallai and colleagues described this type of aberrant spermatogenesis in five globular springtail families, namely Dicyrtomidae (Dallai et al., 1999), Sminthuridae (Dallai et al., 2000), Bourletiellidae (Dallai et al., 2001), Sminthurididae and Katiannidae (Dallai et al., 2004). This is likely the ancestral state of the Symphypleona order. Hence, it is clear that one full haploid set of chromosomes gets eliminated during development (X chromosomes) and spermatogenesis (autosomes) of males. However, it remains unclear whether the chromosome elimination is random during meiosis or systematically dependent on the parental origin (e.g. PGE).

**Figure 2:**
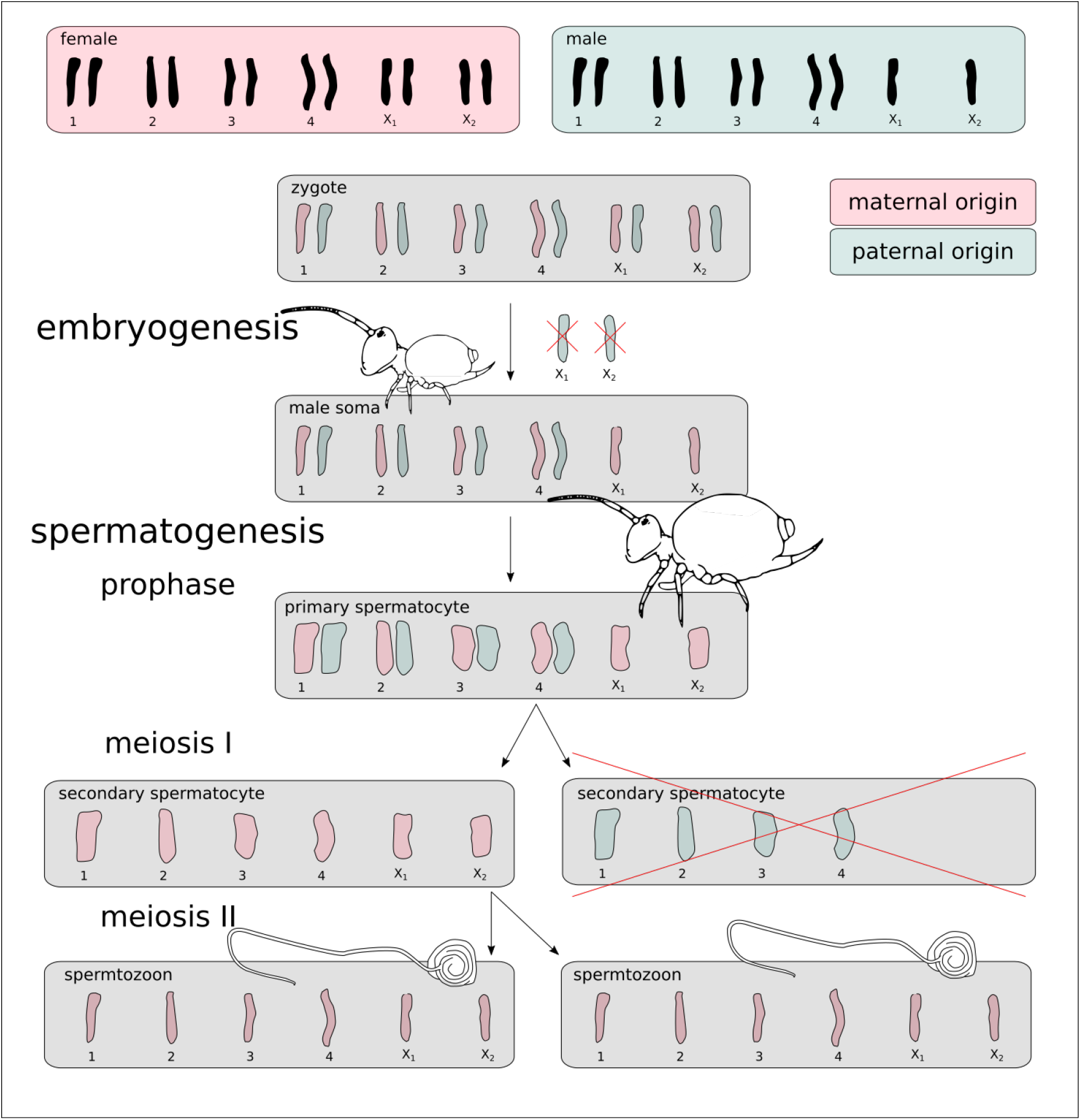
Scheme of the globular springtail PGE model. Male springtail zygotes are initially diploid for all chromosomes. One copy of chromosomes X_1_ and X_2_ is excluded during early embryogenesis. Adult males then generate a half of their secondary spermatocytes with the remaining X chromosomes, and a half without X_1_ and X_2_ that degenerates immediately. The scheme and cartoonized shapes of chromosomes are based on (Dallai et al., 2000). Note the spermatozoon “tail” is not a flagellum, as flagella are densely coiled, see (Dallai et al., 2009) for details. The chromosome movements are well documented (Dallai et al., 2000), but the paternal genome elimination model (the parent-of-origin colouration) is hypothetical, and tested in this study.

There is no distinct name for the putative sex chromosome constellation in globular springtails. It is best described as PGE X0, although the absence of X chromosomes in males is not the primary sex determination. Other springtail orders in contrast have regular meiosis (Dallai et al., 1999) and X0 or XY sex determination (Núñez, 1962; Hemmer, 1990).

We investigated possible approaches to confirm PGE in globular springtails. First, we considered conducting genetic crosses of *Allacma fusca*, a relatively common and large globular springtail commonly found in woodland areas across Europe. However, wild-caught globular springtails are hard to maintain in lab conditions. Alternatively, genotyping a male and its sperm can at least inform us if all sperm contain a single haplotype only, presumably the maternal one. While investigating methods to efficiently sequence male sperm, we discovered male bodies contain a large fraction of sperm (27.5 - 38.4% of cells) and sequencing whole bodies seems to be the most efficient way to sequence sperm, although it requires in-silico analysis to separate the effect of somatic and germline genomes in the sequencing library. We developed and benchmarked a model testing for a mixture of tissues with different karyotypes in a single sequencing library. With our approach we demonstrated that a set of autosomes co-segregate with the X_1_ and X_2_ chromosomes, strongly suggesting uniparental inheritance in *Allacma fusca* males.

## Materials and Methods

### Springtails collected and sequenced

We used an assembly (GCA_910591605.1) and sequencing reads (sample accession ERS6488033) we previously generated for a male *Allacma fusca* individual (Anderson et al., 2020). We also collected 12 additional *A. fusca* samples for resequencing. The sex of individual samples was determined from the modality of sequencing coverage and revealed 11 of 12 resequenced samples were females (**SM Figure 1**). The resequenced male individual was sampled at Blackford Hill (sample id BH3-2, ERS6377982), Edinburgh, Scotland (55.924039, -3.196509). We isolated the DNA using Qiagen DNeasy Blood and tissue kit extraction protocol and sequenced using the Illumina HiSeq platform. The standard adapters and low quality bases were trimmed using skewer `v0.2.2` with options `-m pe -n -q 26 -l 21` (Jiang et al., 2014). We used both the male and all female libraries to identify X-linked scaffolds. Although the reference genome is fragmented, we generated reliable chromosomal assignments for 170.6 Mbp, representing 40.1% of the assembly span. In total, 77.9 Mbp of scaffolds are X-linked, while 92.7 Mbp are autosomal (**SM Text 1**).

All analyses were also performed on the genome of an outgroup species *Orchesella cincta* (GCA_001718145.1, Faddeeva-Vakhrusheva et al., 2016). Both male *O. cinta* resequencing data (ERS7711323) and chromosomal assignments were taken from (Anderson et al., 2020). *Orchesella cincta* is a distantly related springtail with X0 sex determination (Hemmer, 1990) and therefore ideal as a negative control for this study.

### The coverage of unevenly spaced coverage peaks

All sequencing libraries were initially subjected to quality control using kmer spectra analysis. This analysis allows a visualisation and estimate of basic genomic properties without needing a reference genome. We calculated the k-mer coverage histogram with k = 21 using KMC3 (Kokot et al., 2017) and fit a genome model using GenomeScope 2.0 (Ranallo-Benavidez et al., 2020).

In sequencing libraries of a tissue with AAX0 karyotype, the autosomes are expected to have exactly twice the coverage of X chromosomes (i.e. the library has evenly spaced peaks), which is also the expected model of GenomeScope. This would be expected in the soma of male *A. fusca*, as the X_1_ and X_2_ are not homologous chromosomes. However, k-mer coverages displayed unevenly spaced 1n and 2n peaks for the reference *A. fusca* male (**SM Figure 2A**), while the *O. cincta* male showed evenly spaced 1n and 2n peaks (**SM Figure 2E**).

To estimate the 1n and 2n coverage independently, we created a more general model based on similar principles to GenomeScope using non-linear least squares. The adjusted model allowed us to estimate monoploid (1n) and diploid (2n) sequencing coverage for all the sequenced samples. We specifically estimate the ratio between the two coverage peaks and diploid coverage. This formulation of the model allows us to use an asymptotic confidence interval to determine if the coverage ratio of the monoploid and diploid peaks deviates from naively expected 1:2 ratio.

### Two tissue model

The unevenly spaced coverage peaks of X chromosomes and autosomes imply the sequencing library contained tissues with various ploidies in *A. fusca.* A simple model that can explain the pattern is a two tissue mixture model - a mixture of a tissue with an X to autosome ratio of 1:2 (e.g. male soma) and a tissue with an X to autosome ratio of 1:1 (e.g. secondary spermatocytes or sperms). Using the coverage estimates of X-chromosome and autosome peaks, we can estimate the relative contribution of the two tissue types to the sequencing library (**SM Figure 3A**) and the fraction of the two tissues. Assuming the 1:1 tissue is haploid, the relative fraction of that tissue (*f_h_*) in the sequencing library is

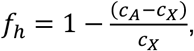

where *c_A_* is the coverage of the autosomal peak, and *c_X_* is the coverage of the X chromosome peak. We estimated the fraction of the haploid tissue using both k-mer coverage estimates (described above) and mapping coverage (**SM Text 2**). While the estimates from mapping coverage have lower sampling variance, they rely on a well assembled reference genome and therefore are less suitable for non-model species. For detailed explanation of these different types of coverages see (**SM Text 3**).

The only described tissue with 1:1 X to autosome ratio in adult male globular springtails are primary and secondary spermatocytes, spermatids, and spermatozoa (Dallai et al., 2000). Hence, it is probably safe to assume this is the tissue that is causing the relative mapping coverage shift illustrated in **SM Figure 3B** (for alternative unsupported hypotheses tested to explain the 1n mapping coverage shift, see **SM Text 4** and **SM Figure 4**).

We validated the power available to estimate the proportion of sperm in the body using the relative positions of the two coverage peaks (two tissue model) using a power analysis. We simulated genomes with various X chromosome sizes, levels of heterozygosity, sequencing coverage and fraction of sperm present (see **SM Text 5** for details).

### Testing PGE

Sequencing a mixture of sperm and soma allows us to test the suggestion that globular springtails reproduce by PGE (Dallai et al., 2000). The PGE inheritance model (**Figure 2**) predicts that sperm contain only the maternally inherited haploid set of chromosomes (A_m_X_m_). As all the autosomes present in the haploid sperm are of maternal origin, all the heterozygous autosomal loci should display a higher coverage support of maternal alleles compared to paternal. The k-mer coverage we used in the two tissue model does not directly translate to allele coverage (see **SM Text 2**). To calculate the exact expected coverages of maternal and maternal alleles, we used the estimated fraction of sperm (*f_h_*, estimated by the two tissue model) and mean allele coverages of homozygous autosomal variants (*c_AA_*). The maternal and paternal coverage expectations are

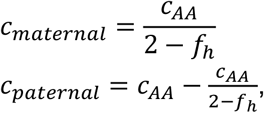

In the ideal case, we would like to compare the expectations to coverage supports of phased haplotypes, which is unfeasible with fragmented reference genomes and short read libraries. Instead, we separated the alleles of heterozygous autosomal variants to the “major” and “minor” alleles - representing the variants with higher and lower coverage support respectively. Under the PGE model the maternal and paternal alleles are expected to have vastly different coverage support, therefore the “major” alleles will be vastly of maternal origin, while the “minor” alleles will be vastly paternal. The fraction of possible misassigned variants was explored through modelling of sequencing coverages using negative binomial distributions with parameters estimated from expected sequencing coverages.

Furthermore, under the PGE model, the distribution of maternal allele coverage depths is expected to resemble the distribution of X-chromosome allele coverage depths. Due to a small fraction of misassigned alleles in males (as explained in the previous paragraph) the match is not expected to be exactly perfect. The expected levels of imperfect match were also estimated via the same set of simulated coverages.

We performed the same analysis on the genome of male *O. cincta* and two *A. fusca* females. The two females only show the decomposition of autosomal heterozygous alleles in the case of frequent misassignment of maternal and paternal alleles (as they are generated from the same coverage distribution, **SM Figure 5**). The *O. cinca* male further allows the same comparison of decomposed allele coverages to the distribution of coverage of alleles found on the X chromosome.

To test the PGE model we mapped trimmed sequencing libraries to the softmasked reference genomes of *Allacma fusca* (`GCA_910591605.1`) and *Orchesella cincta* respectively (`GCA_001718145.1`). The reads were mapped using bowtie2 with the parameter `--very-sensitive-local` (Langmead & Salzberg, 2012). Before calling variants we marked duplicates in the mapping files using picard MarkDuplicates (Picard toolkit, 2019) and called variants using freebayes `v1.3.2-dirty` (Garrison & Marth, 2012) with stringent input base and mapping quality filters as well as required minimal allele coverage (`--standard-filters--min-coverage 5`), but we relaxed the priors of Hardy–Weinberg proportions as they might not be met in a PGE population (`--hwe-priors-off`), while assuming diploidy (`-p 2`). The raw variant calls were subsequently filtered for high quality variants (`-f “QUAL > 20”`) only using vcffilter from the vcflib library version `1.0.0_rc3` (Garrison et al., 2021) and sorted to autosomal and X-linked using a custom python script. The variants sorted to chromosomes were plotted using R scripts.

### An alternative estimate of the relative fraction of the haploid tissue

In the section above we showed how the fraction of haploid tissue (*fh*) can be estimated using estimates of X chromosome (1n) and autosomal (2n) coverages. This formula works regardless of the chosen type of coverage, therefore we applied it both to the k-mer coverages presented in the main text and mapping coverages (**SM Text 2**).

Assuming the PGE model, however, we can also estimate the fraction of sperm in the male body from the minor allele frequency. As all the sperm is expected to contain only maternally inherited autosomes, the expected proportion *p_p_* of paternal (green shaded in **Figure 2**) autosomes over all the body’s cells is 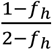. The expected allele coverage ratio (site frequency) of the paternal state is *p_p_*, this is the minority state when *f_h_* > 0, and

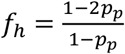

which allows us to estimate the relative fraction of haploid tissue (sperm) from the estimated allele coverage ratios. This approach can be applied to heterozygous SNP calls (**SM Figure 6**) and even to raw pileups.

We explored the raw pileups to avoid the lack of power and any other potential biases introduced via SNP calling. For the two *Almaca* males we counted sequence states aligned under the `GCA_910591605.1` reference using samtools mpileup converted to matrix form with Popoolation2 mpileup2sync (Kofler et al., 2011). After filtering out scaffolds with evidence of copy-number variation between the males (**SM Figure 7**), we calculated minor frequencies *p_p_* for all genomic positions with at least two states in the pileup. Then we examine the distribution of variant sites by minor allele frequency for both males. See **SM Text 6** for details.

## Results

The analysis of trimmed sequencing libraries of the two *A. fusca* male individuals revealed that both have unexpected relative k-mer coverages of the monoploid k-mers (X chromosomes and heterozygous loci) compared to the diploid k-mers (autosomes). In both males the 1n coverage estimates were more than half of the diploid coverage estimate. We estimated the coverage ratio of the X-chromosome to autosome in the BH3-2 male to be 0.607, 0.95 asymptotic CI [0.582, 0.633]), significantly deviating from the 1:2 ratio. A remarkably similar coverage ratio of the two unevenly spaced peaks was observed in the reference *A. fusca* male: 0.58, 0.95 asymptotic CI: [0.576, 0.584]., The coverage ratio in *A. fusca* males was in a strong contrast with a male sequencing library of a non-PGE species *O. cincta*, where the two coverage peaks were nearly perfectly spaced, with the coverage ratio 0.501, 0.95 assymptic CI: [0.499, 0.503] as expected for an XO species (**SM Figure 2**).

Using the coverage estimates and the two tissues mixture model (see **Methods** and **SM Figure 5**) we calculated the fraction of sperm cells in male *A. fusca* to be 27.5% (the reference male) and 35.3% (BH3-2, **Figure 3**). These estimates were comparable to the estimates using mapping coverage instead of the k-mer based estimate (**SM Text 2** and **SM Table 3**). For comparison, the fraction of sperm that would be estimated in *O. cincta* male if we have (wrongly) assumed PGE model is 0.6%.

**Figure 3:**
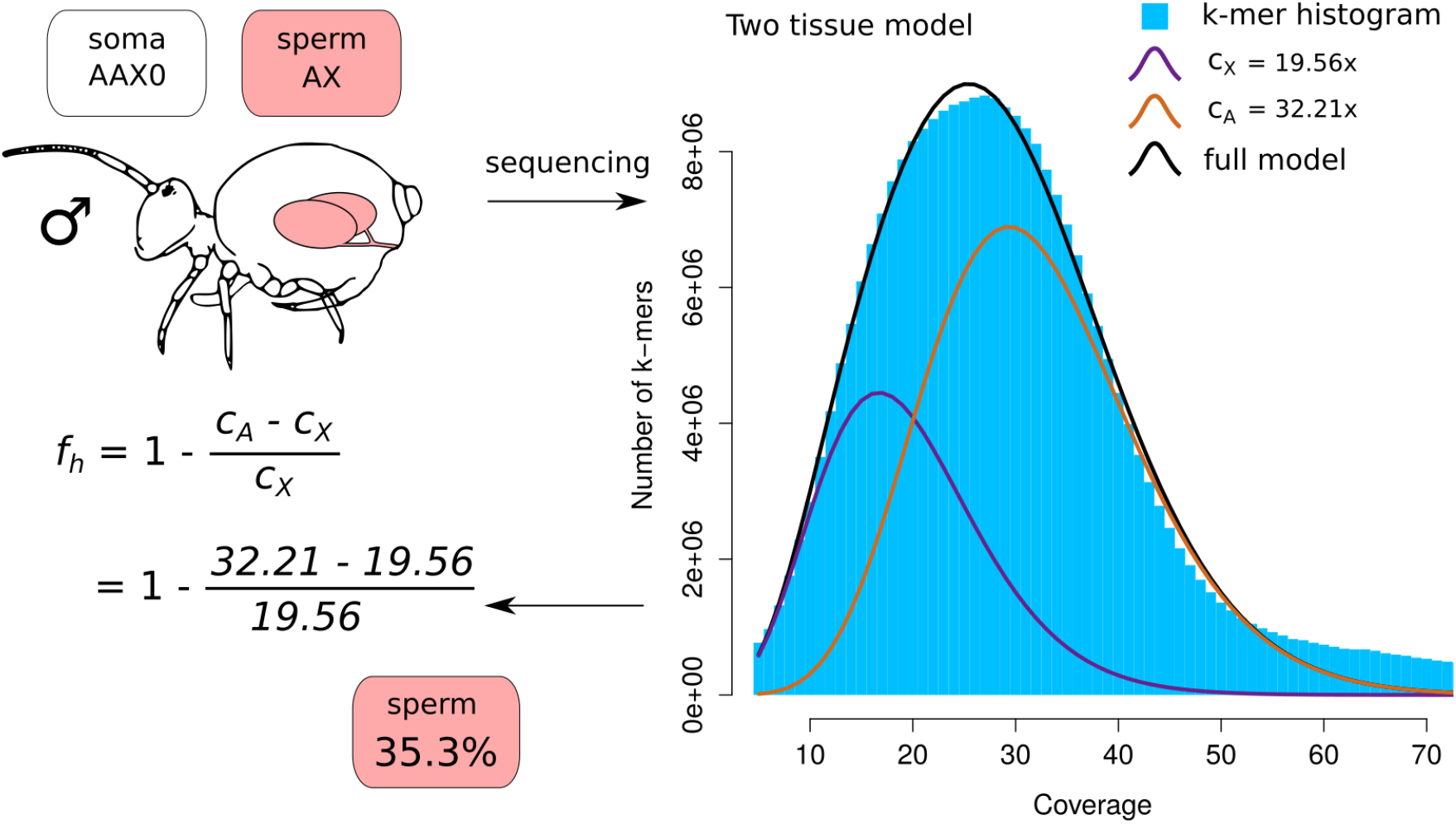
Overview of how expected coverages potentially explain the shift of coverage peaks. PGE is expected to cause the shift of coverage peaks due to a significant proportion of sperm in the body, as indicated in cartoons (explained in greater detail in **SM Figure 3**). The table contains the expected paternal and maternal coverages of autosomes and X-chromosomes for the male resequencing individual BH3-2.

Using the estimated fraction of sperm in *A. fusca* males we calculated the expected coverages of paternal and maternal autosomal alleles for the PGE model (**Figure 2**). For the BH3-2 individual the estimated allele coverage expectations are 11.29x for paternal and 17.44x for maternal autosomes and X chromosomes respectively.

The expected coverages of maternal and paternal autosomes and X chromosomes were compared with the distribution of allelic coverages of variants on autosomes and X chromosomes. After quality filtering we identified 28,070 and 235,301 heterozygous variants anchored to chromosomes in the reference and BH3-2 individuals respectively (**SM Table 2**). The extremely low heterozygosity of the reference male reduces the power to use the sample for testing the PGE hypothesis and is discussed in **SM Text 7**. Of the BH3-2 anchored heterozygous variants, 227,570 were located on autosomal scaffolds, while only 7,731 heterozygous variants were on X-linked scaffolds, indicating low levels of false positives among variant calls (less than 100 false positives per 1Mbp). On the other hand, we identified 60,999 homozygous variant calls on the X linked scaffolds that were used for the comparison with allele coverages of the autosomal variants. The coverage supports of *A. fusca* male were contrasted to 1,959,258 heterozygous autosomal variants and 400,001 homozygous X-linked variants in the outgroup species *O. cincta* (non-PGE springtail).

We decomposed the male heterozygous autosomal variants in both samples to the “major” and “minor” alleles - representing the variants with higher and lower coverage support respectively. Given the PGE model and the estimated fraction of sperm, the mean coverage of maternal variants (17.44x in BH3-2) is expected to be higher compared to the coverage of paternal variants (11.29x), hence although it is possible some of the paternal variants will be by chance higher, this will affect only a very small fraction of the variants. On the other hand, applying the same decomposition of heterozygous variants to “major” and “minor” in non-PGE species leads to ~50% of misassigned variants (by definition). To demonstrate the effect of misassigning variants by coverage, we simulated the coverage of maternal and paternal alleles under the PGE model (**Figure 4A**) and regular X0 species (**Figure 4B**). In both cases the black distribution in the background represents the background distribution for the maternal variants. In the real data, we used the homozygous variants located on the X chromosomes to estimate the coverage distribution of monoploid maternal alleles. Under the PGE model, we expect it to roughly overlap with the “major” variant coverage peak (**Figure 4A**), contrasting to the non-PGE model where the expected distribution will be exactly in the middle of the “major” and “minor” coverage peaks (**Figure 4B**).

**Figure 4:**
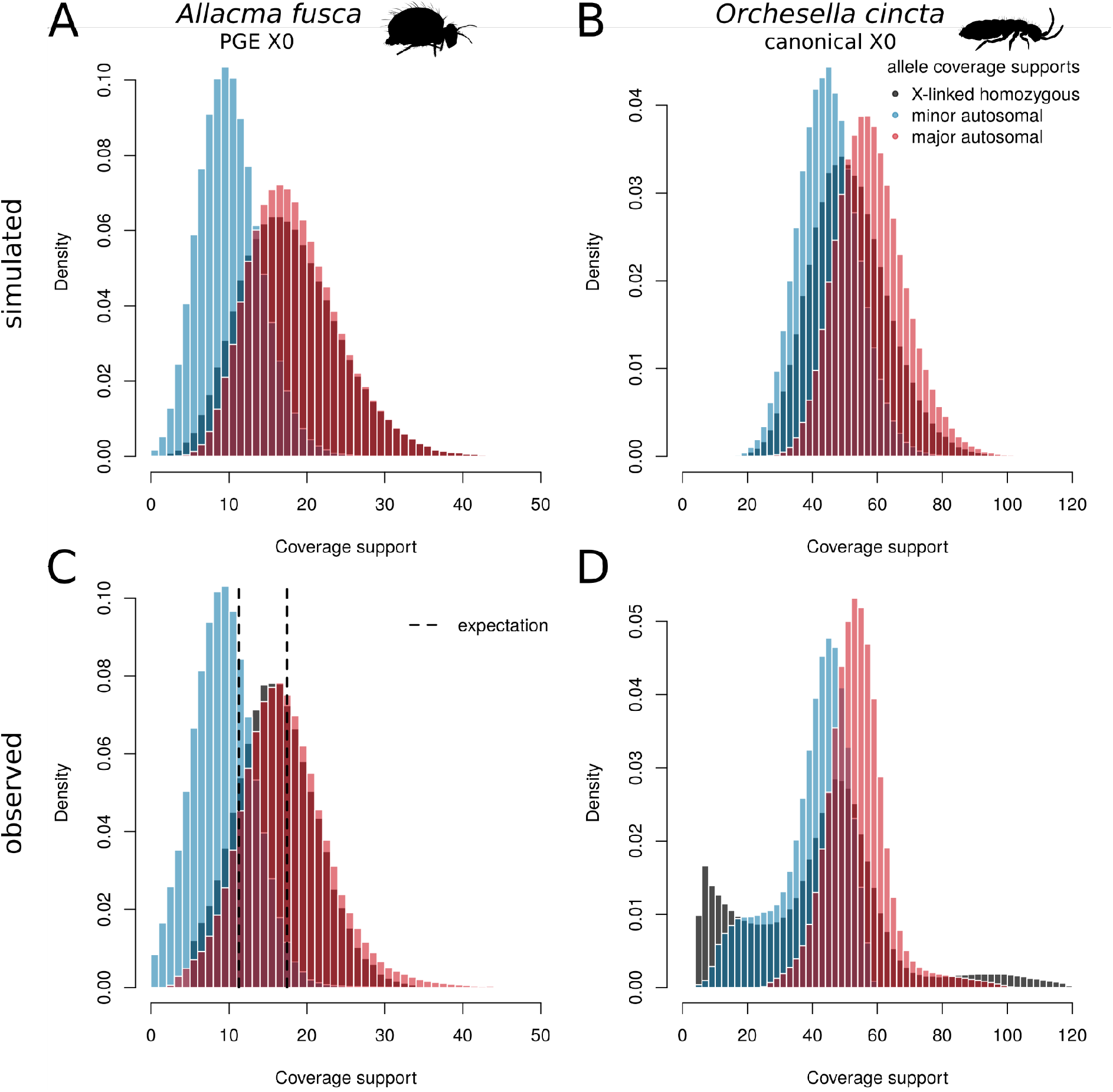
Decomposed heterozygous allele coverage supports. Coverage supports of the two alleles of heterozygous sites are decomposed to those with higher coverage (“major”, in red) and lower coverage (“minor”, in blue). These are compared to coverage supports of homozygous X-linked variants. Panels **A** and **B** are simulated allele coverages for a PGE X0 system and non-PGE X0 system using negative binomial with means corresponding to means of empirical data and size parameter 15. In PGE species (**A**), major alleles are almost all maternal alleles and show similar coverage distributions to homozygous X-linked alleles (maternal haploid). In canonical X0 system (**B**) the decomposition also leads to bimodal distribution, however, the X-linked allele has an intermediate coverage peak in between of the two autosomal distributions. The observed coverage distributions in *Allacma fusca* (**C**) strongly support the PGE model. The major allele coverage distribution closely resembles the distribution of homozygous X-linked alleles as well as matching the expected coverage calculated from the 1n coverage shift (**Figure 3**). In contrast *Orchesella cincta* (**D**), a species with regular meiosis and X0 sex determination, shows patterns consistent with the expected properties of canonical sexual X0 sex determination species with X-linked coverage support intermediate of the decomposed autosomal coverage supports.

We confirm the coverage supports of “major” and “minor” autosomal variants in *A. fusca* male BH3-2 are close to the expected coverages generated using the two tissue model (**Figure 4C**). The fit is not perfect, probably due to misassigned alleles. Furthermore, the distribution of “major” autosomal variants closely resembles the distribution of homozygous X-linked variants, with similar levels of disagreement compared to the simulated data (**Figure 4A**). Both comparisons together provide strong support for the PGE model in *Allacma fusca.* The analysis of *A. fusca* shows a clearly different pattern to *O. cincta*, a springtail with standard meiosis. The decomposed coverage supports display largely overlapping distributions and the coverage distribution of X-linked variants is nearly located intermediate between the peaks of “major” and “minor” allele coverages (**Figure 4D**), as predicted by the non-PGE model (**Figure 4B**). Note that the first coverage peak of X-linked variants displays spurious and unexpected coverages, which according to the genome profiling (**SM Figure 2E**) should be considered false positives.

Additionally, we utilised an analysis of raw pileups to create an independent estimate of the fraction of the haploid tissue *f_h_* from the estimated minor allele frequency of all genomic positions with two states located on scaffolds with no signs of copy number variation (See **SM Text 6** and **SM Figure 7**). We used genomic positions with two nucleotides with coverage >1 mapped to it. This approach showed a higher abundance of these bistates around coverage ratios 0.397 in Afus1 and 0.406 in BH3-2 (**SM Figure 8**), indicating that even the reference male shows some detectable heterozygous states, but with much noisier signal compared to BH3-2. The estimated fraction of sperm in the bodies from the paternal allele frequency *pp* are 33.96% for Afus1 and 31.39% for BH3-2 individuals respectively. Overall both types of estimates of fractions of sperm (based on shift of the X chromosome coverage peak, and minor allele frequency) showed relatively consistent levels (**SM Table 3**)

## Discussion

We estimated that a large proportion of a male adult *A. fusca* body (>27.5%) consists of secondary spermatocytes, spermatids or mature sperm (from now on collectively referred to as sperm). Although the estimated fraction is relatively high, it is in agreement with the high production of spermatophores by *Allacma fusca* (Dallai et al., 2009) and the estimate does not surpass that of other invertebrates. *Caenorhabditis elegans* can carry around 2000 germ cells, while their soma consists of precisely 959 cells (Sulston & Horvitz, 1977). Germ cells therefore represent ~67% of *C. elegans* cell count. A similar case is found among arthropods: Up to 75% of body cells in *Daphnia* males are sperm (Dufresne et al., 2019). It is important to note we specifically discuss the fraction of cells, not the biomass, as sperm can be substantially smaller compared to other cell types in the body.

We performed a power analysis (**SM Text 5**) to describe biological conditions for which such analysis would be possible assuming the model shown on **Figure 2**. We revealed that for X chromosomes spanning more than 10% of the genome we managed to detect a significant deviation of 1:2 coverage ratio of the two coverage peaks in 132 out of 144 cases (**SM Figure 10**). In general, those with greater coverage converged more often and levels of heterozygosity had a surprisingly small effect on convergence of the two tissue models. We found the fraction of sperm is systematically underestimated using our technique and therefore the results are likely a conservative estimate of the real fraction of the haploid tissue in globular springtails. Finally, we demonstrated that the two tissue model can be fully automated to scan for presence of multiple karyotypes in a library for the majority of parameter combinations. Therefore it might be useful for naive scans in the large number of genomes currently sequenced without any previous cytological studies (The Darwin Tree of Life Project Consortium, 2022).

Taking advantage of the high sperm fraction, we demonstrated that all the sperm have exactly the same genotype which conclusively implies co-segregation of full chromosomal sets under the absence of recombination in this globular springtail. This conclusion was also supported by analyses of read pileups (**SM Text 6** and **SM Figure 8**). The analyses of pileups further revealed that the reference sample indeed also shows uneven coverage ratios of heterozygous alleles, although this signal was much weaker compared to the BH3-2 individual. We propose the reference individual could have reduced heterozygosity due to local inbreeding of the population that was sampled. Altogether, all results are in agreement with the PGE model (**Figure 2**) that has been previously proposed (Dallai et al., 2000).

We have shown a set of chromosomes is eliminated, but not whether the eliminated set is maternal or paternal. To provide definitive proof of PGE we would have to genotype both parents of a male as well as its sperm. Hypothetically, the eliminated chromosomes could be maternal. However, the elimination of maternal chromosomes during spermatogenesis has only ever been observed in a rare form of androgenesis (Schwander & Oldroyd, 2016), a reproductive system in which males fertilize a female of a closely related sexual strain and cause elimination of the maternal genome as found in freshwater clam *Corbicula leana* (Komaru et al., 1998) or in combination with hybridogenesis in Australian carp gudgeons (Majtánová et al., 2021). However, this form of androgenesis requires a co-existence of lineages with canonical sexual reproduction with male androgenetic lineages, which is extremely unlikely in the case of globular springtails as aberrant spermatogenesis seems to be already present in the common ancestor of globular springtails (Dallai et al., 1999, 2000, 2001, 2004). Paternal genome elimination on the other hand is a mode of reproduction that is conserved in at least six large clades (**Figure 1**) and although with our data we also cannot completely exclude the possibility that the non-random chromosome elimination is associated with a different, as yet undescribed, evolutionary phenomena, paternal genome elimination is the only explanation compatible with known biology.

In particular, globular springtail reproduction most closely resembles the reproductive cycle of two dipteran families that also eliminate paternal chromosomes both in early spermatogenesis (which we call X chromosomes in these species) and during spermatogenesis (Metz, 1938; Gerbi, 1986). In both these two families females are frequently monogenic - each female produces broods of a single sex only (Metz, 1931). So far this has not been tested in globular springtails, probably because they are both difficult to cultivate and show very little sexual dimorphism. Finally, the third genomic peculiarity found in both PGE fly families - they carry germ-line restricted chromosomes (Metz, 1938; Hodson et al., 2021), is a feature that is not shared with globular springtails as no differences between germ-line and soma karyotypes have been reported other than the aberrant spermatogenesis discussed in detail above.

Although we have tested this hypothesis in only a single globular springtail species *Allacma fusca*, the same type of aberrant spermatogenesis was demonstrated in seven species of five different families (**SM Table 1**, Dallai et al., 1999, 2000, 2001, 2004). The most parsimonious explanation of the aberrant spermatogenesis in all the examined species is that PGE is the ancestral feature of globular springtails. Although we expect most of the globular springtails to retain this type of reproduction, there are multiple transitions to parthenogenetic reproduction (reviewed in Chernova et al., 2010). Other PGE clades usually show high conservation of this reproduction mode (Brown, 1965; Gerbi, 1986; Ross et al., 2012), the only known exception is found in lice. The human body louse seems to show a partial reversal to a non-PGE sexual type of reproduction (McMeniman & Barker, 2005; de la Filia et al., 2018). Whether or not any globular springtail species have reverted to a more canonical type of reproduction is however an open question for further research.

Our study strongly suggests that globular springtails are the oldest and most species-rich clade reproducing via PGE. With 15,600 species estimated worldwide (Porco et al., 2014) globular springtails are a great clade to study the long term consequences of coping with PGE over hundreds of millions of years of evolution (Leo et al., 2019). This unusual mode of inheritance is likely to profoundly influence their evolutionary history. Recent theory suggests that haplodiploidy and PGE affect the evolution of reproductive isolation and increase diversification rates because of a generation lag of hybrid males that can be produced only if the mother is a hybrid already (Patten et al., 2015; Lohse & Ross, 2015). Springtails provide a great opportunity to test this theory as three of four springtail orders are species rich and allow us to estimate rates of diversification. Paternal genome elimination also affects the dynamics of sexual conflict as shown in recently developed models (Hitchcock et al., 2021; Klein et al., 2021). Notably, it changes the relative role of X chromosomes and autosomes. Under PGE both X chromosomes and autosomes show bias in transmission between generations and sex alternation (see Klein et al., 2021 for details), however, X chromosomes in globular springtails are also subjected to haploid selection in males. Unlike in species with normal diploid reproduction, the dominance of male beneficial alleles is the only factor that determines if they are more likely to get fixed on X chromosome (for recessive alleles) or anywhere in the genome (for dominant alleles) (Klein et al., 2021). Therefore comparing the levels of sexual antagonism on X chromosomes and autosomes in globular springtails will allow the effect of dominance in sexual selection to be quantified, which has been a central question of sex chromosome evolution.

## Data, script and code availability

All the raw data are deposited in ENA/SRA under accession PRJEB44694. All code and processed output is archived at doi.org/10.5281/zenodo.6645407.

## Supplementary material

**Supplementary texts, figures and tables** and

**Supplementary Table 1** and available online doi.org/10.1101/2021.11.12.468426.

## Acknowledgements

First of all, we would like to thank Tanja Schwander for the invaluable suggestion of sperm being the cause of disrupted coverage ratio of autosomes and X chromosomes in males, Hannes Becher for designing large part of the power analysis workflow and Saudamini Venkatesan for help with statistics. We would also like to thank Konrad Lohse, Andrew Mongue and the rest of Ross lab members for useful discussions and comments on earlier versions of this manuscript. Wytham Woods, City of Edinburgh Council, and Friends of the Hermitage of Braid for permission to collect springtails. Version 5 of this preprint has been peer-reviewed and recommended by Peer Community In Evolutionary Biology (https://doi.org/10.24072/pci.evolbiol.100142).

## Funding

CH was supported by NSERC and the Darwin Trust of Edinburgh for postgraduate financial support. LR would like to acknowledge funding from the European Research Council Starting Grant (PGErepo) and from the Dorothy Hodgkin Fellowship DHF\R1\180120. SJEB was supported by MOZOOLEC: Mobility project to support international research collaboration in zoology and ecology, grant number CZ.02.2.69/0.0/0.0/18_053/0017792.

## Conflict of interest disclosure

The authors declare they have no conflict of interest relating to the content of this article.

## Notes

https://github.com/RossLab/genomic-evidence-of-PGE-in-globular-springtails

### Competing Interest Statement

The authors have declared no competing interest.

### Summary of Updates

Fixed metadata for PCI recommendation!

