## Supplementary texts, figures and tables for "Genomic evidence of paternal genome elimination in the globular springtail *Allacma fusca*"

### SM Text 1: X chromosome assignments

The X chromosomes were annotated on GCA_910591605.1 assembly, using eleven females and two males (all samples accessible via project with PRJEB44694 EBI accession). The reads were mapped in the same way as described in methods. We extracted per base coverage using `samtools depth` (Li et al., 2009) and a custom script to generate a table of per scaffold coverages for all thirteen samples. We kept only scaffolds longer than 10,000 bases for chromosomal assignments, as shorter scaffolds show high variation in coverages.

We used the same approach based on kernel smoothing to estimate the 1n and 2n mapping coverages in all the samples (both male and female). The 2n coverage estimates were used afterwards for normalization of per scaffold coverages. For each scaffold we calculated median and variance of normalized coverages among male and female samples. Their $\log_2$ ratio was then used to determine scaffolds considered as X-linked or autosomal respectively.

In these analyses, usually X-linked staffolds show male to female $\log_2$ ratio around -1, while autosomal peak is expected around 0. A commonly used threshold to annotate scaffolds is > -0.5 for autosomal and <-0.5 for X-linked scaffolds. However, male *A. fusca* sequencing library is a mixture of two tissues with different karyotypes (**SM Figure 3**). As a consequence, $\log_2$ normalised male to female coverage ratio of X chromosomes will be shifted. Hence, we used kernel smoothing to determine the local minimum between the two coverage peaks, which is -0.4205 and used that as the threshold for separating autosomal and X-linked scaffolds.

Finally, we used female samples to determine which scaffolds show spuriously high variation in coverages, possibly due to structural variants and polymorphic transposon insertions. We assigned to chromosomes only the scaffolds with variance among female samples <100. This threshold was chosen manually after exploration of the distribution of coverage variations.

With this procedure we assigned in total 170.6 Mbp of scaffolds to chromosomes, corresponding to 40.1% of the assembly span. In total, 77.9 Mbp of scaffolds are X-linked, while 92.7 Mbp are autosomal. These scaffolds were used in all the subsequent analyses.

### SM Text 2: Mapping coverages

The main approach we utilised to estimate coverage of X chromosomes (1n) and autosomes (2n) was used on reference-free decomposition of sequencing reads into kmers. While the k-mer technique has an advantage of being reference-free, it requires sufficient coverage and furthermore the X-chromosome signal is co-founded with kmers originating from heterozygous loci. Alternatively, we can estimate these coverages by mapping reads to a reference genome.

We used the same mapping files (.bam) as for the SNP calling. We used samtools depth to extract per base coverage (Li et al., 2009), and calculated per scaffold mean coverage. Then we estimated 1n and 2n coverage peaks using kernel smoothing with kernel width chosen by Sheather and Jones method (`bw = "SJ"`) (Sheather & Jones, 1991) while weighted by scaffold length (`weights = scf_tab$len / sum(scf_tab$len)`).

This method resulted also in both *A. fusca* males showing distinctively uneven peaks (**SM Figure 2B** and **D**). *O. cincta*, however, also showed a substantial deviation from the 1:2 coverage ratio. Applying the two tissue model using 1n and 2n coverage estimates from mapped reads (53.3x and 101.1x respective) results in an estimate of 10.2% of sperm with a different karyotype. This is very likely caused by several scaffolds in the *O. cincta* are chimeric - partially containing autosomal and partially X-chromosome sequences. As a consequence, these scaffolds have coverage somewhere in between 1n and 2n peak and cause them to be “smoothed” to each other than they biologically should have been.

However, note that the mapping approach is very robust when a correct reference genome is provided, as demonstrated in power analysis (**SM Figure 10**). We made no assembly errors to the simulated reference genomes therefore this downside of the mapping approach was not part of the analysis.

### SM Text 3: Different types of sequencing coverages used in this manuscript

Sequencing coverage is the mean number of times every position in a genome is represented in reads. Sequencing coverage is usually estimated by dividing the total sequencing yield by the haploid genome size. However, in many cases, for genome analyses of non-model organisms the level of contamination in sequencing libraries; sequencing errors; and genome size is unknown. This can make estimating sequencing coverage challenging. Hence there are multiple other ways to measure and estimate sequencing coverage. These different measures have different properties and are used for different purposes. **K-mer coverage** is the mean number of occurrences of each unique continuous genomic sequence of length k in reads. K-mer decomposition is independent of a reference genome and the coverage is estimated by fitting a model to k-mer coverage histogram (Ranallo-Benavidez et al., 2020). This reference-free technique is well suited to observing raw signals from data unbiased by complicated procedures such as genome assembly. **Mapping coverage** is the mean number of reads mapping to each position of a haploid reference genome. It is dependent on the quality of the reference and quality of mapping. The main advantage is that reads of heterozygous sites as well as infrequent sequencing errors still typically map to the same position on the reference. Hence, this coverage is suited the best to estimate ploidy of each reference genomic region. **Allelic coverage** is also derived from sequencing reads mapped to the genome. However, the mapped reads usually have PCR duplicates marked and are subsequently used for calling variants. The coverage is then the number of non-duplicated reads supporting individual alleles. Sequencing errors do not contribute to this coverage.

Note that the k-mer coverage ($C_k$) and mapping coverage ($C_m$) have different coverages for the same sequencing dataset. Assuming no sequencing one can be converted to the other by a simple approximation

$$C_k \approx C_m \frac{R-k+1}{R}$$

Where $k$is the length of k-mer, $R$ is the length of reads. To consider errors we need to multiply mapping coverage by fraction of correct k-mers in the dataset

$$C_k \approx C_m \frac{R-k+1}{R} (1-e)^k$$

Where $e$ is the sequencing error rate. However, in practice there are usually too many issues - no haploid reference is perfect, sequencing errors are not uniformly distributed along reads, and the mapping process is also dependent on many assumptions. Hence in practice the two measures need to be calculated independently.

### SM Text 4: Alternative unsupported explanations of 1n coverage shift peak

We generated two additional models that could explain the 1n coverage shift: *imperfect chromosome elimination* and *tissue specific endoduplication*. These hypotheses as well assume that we sequence a mixture of two tissues: the soma and a tissue that is diploid also for the two X chromosomes (**SM Figure 4**). The *imperfect chromosome elimination* hypothesis assumes that the elimination process during early development is imperfect and some of the cellular lineages simply retain two copies of the X chromosomes ($A_mA_pX_mX_p$). The *tissue specific endoduplication* hypothesis assumes endoduplication of the two X chromosomes in some specific tissue, presumably to allow higher expression levels of genes located at these chromosomes. The endoduplicated tissue would be expected to have then two, but the same copies of X chromosomes ($A_mA_pX_mX_m$).

Both these hypotheses can be rejected as there are coverage differences in coverages of autosomal heterozygous loci as shown on **Figure 3**. Furthermore, we have observed only a very few heterozygous loci on male X chromosomes (6,599 and 7,731, see **SM Table 2**), suggesting these are false positive calls rather than real heterozygous loci, which is a second line of evidence refuting the *imperfect chromosome elimination* hypothesis as no paternal alleles were discovered with lower coverage supports. The *tissue specific endoduplication* would also cause that the coverage of X-linked alleles would be higher compared to autosomes. This is, however, not the case as the major allele coverage distribution resembles the distribution of X-linked variants (**Figure 3**).

Finally, none of these two alternative explanations have any foundation in cytological studies previously done on globular springtails. In conclusion, other mechanisms causing the 1n shift were considered, but none of the proposed explanations is compatible with observed data.

### SM Text 5: Power analysis

We explored the strength of signal that would be needed to detect a significant distortion of the 1:2 coverage ratio of the 1n and 2n coverage peaks. We explored gradients of the X-linked portion of the genome, heterozygosity, fraction of sperm and sequencing coverage as factors that might contribute to the power of detecting the signal of paternal genome elimination.

This has been done using simulation workflow in following steps:

1. Generate 20 chromosomes, 1 Mbp each. In the simulations 1, 2, 5 or 10 of the chromosomes were X chromosomes and the rest was autosomal (corresponding to 5, 10, 25 and 50% of the genome being X). These chromosomes were made out of randomly selected and then catenated contigs from the *Allacma fusca* reference genome with autosomal or X-chromosome assignment respectively.
2. For each chromosome we run a classical coalescence with recombination model simulation (Hudson's algorithm) using msprime (Kelleher et al., 2016). This approach allows us to mimic a more realistic distribution of heterozygous loci across the genome. We used 5e-7 per base recombination rate and effective population size of 1000 individuals with a gradient of mutational rates so it corresponds to various levels of heterozygosities (0%, 0.01%, 0.1%, 0.5%). Both maternal and paternal haplotypes were created.
3. The simulated read coverage from the templates was dependent on maternal coverage (10x, 15x or 25x) and fraction of sperm (0.0, 0.01, 0.05, 0.1, 0.25, 0.50). For each combination of parameters the corresponding maternal coverage was generated out of the mutated autosomal and X chromosome and paternal coverage from the autosomes only (following PGE model). The reads were simulated 150 bases long with flat 0.01 sequencing error rate using wgsim (https://github.com/lh3/wgsim).
4. The simulated reads were decomposed into the k-mer database and subjected to the same two-tissue analysis as the biological data generated in this study. The fraction of sperm estimated using this approach is called "kmer estimates".
5. Reads were mapped back to the original unmutated template genome (step 1). Then we extracted coverage per each non-overlapping 10,000 genomic window. These coverages were used to estimate 1n and 2n coverages using kernel smoothing in a very similar way as it was done for the biological samples (**SM Text 2**). The coverage estimates were then used to create an alternative estimate of the fraction of the sperm ("mapping estimates")

The major effect was due to the size of the X chromosome. With 5% of genome X-linked, the model frequently did not converge, especially for the low coverage datasets. With 10% of the genome being X-linked nearly all models converged although the 1n and 2n coverage estimates of overlapping peaks was posing a major problem for low coverage datasets, especially for simulations with low levels of sperm. For X chromosome linkage greater than 10% models generally converged, but only for proportion of sperm >10% the model was significantly different to 0. In general we found that various levels of heterozygosity have very little effect on the quality of the estimates. Finally, it seems the estimate of the fraction of sperm is slightly negatively biased, indicating that the genomic estimates are on the conservative side. The full parameter grid was plotted on a single figure **SM Figure 10.**

#### SM Text 6: Fraction of sperm estimated from raw pileups

The two tissue model and the assumption of PGE allows an independent estimate the relative fraction of the haploid tissue of $f_h$based on a similar idea as the allele coverage distribution test of autosomal heterozygous variants, but avoiding the stringent parameters of the SNP calling pipeline.

For male whole-body sequencing in the absence of (a) mutation relative to the gametic sequences entering the zygote, (b) sequencing error and (c) mapping error, the only variant sites when reads are aligned under a reference should be bi-states - heterozygous loci: those where paternal and maternal autosomes differ in their sequence state. As mentioned in methods, the expected mapping coverage (site frequency) of the paternal state is $p_p$, this is the minority state when $f_h$> 0, and $f_h = \frac{1-2p_p}{1-p_p}$therefore we can in principle estimate $f_h$ from the frequency of the minor state over all variant aligned sites, with the caveat: If autosomes contributed by male and female gametes are identical, then there is no power, as we expect zero variant sites (i.e. low $f_h$estimation power in the case of inbreeding). Deviations (a,b,c) from the assumed ideal suggest how $f_h$estimation should proceed. (a) Whole-body sequencing reads will include somatic and germline point mutations, producing apparently variant sites. As long as the two point mutation rates are low relative to $p_p$, it should be straightforward to distinguish point-mutation-variant sites from gamete-variant sites. Further, as long as (b) the (pointwise) sequencing error process is independent from (a) the mutation processes, and its rate low relative to $p_p$, it should remain straightforward to distinguish all (a,b) pointwise-change-variant sites from gamete-variant sites. Mapping error (c) will obscure this clarity. Where reference and assayed individuals differ in motif copy number along the genome (copy number variation, CNV), mapping will overmerge or undermerge reads. Where a motif occurs *less* in the reference, overmerged assayed reads will produce apparent CNV-variant sites. Where a motif occurs *more* in the reference, undermerging of assayed reads can reduce minor state frequency at otherwise gamete-variant sites. When we also allow for somatic and germline CN mutation processes, and their resultant over/under merging during mapping, we may expect the distribution of variant sites by minor frequency to have two peaks: (1) At low minor frequency (pointwise-change-variant); (2) at higher minor frequency (gamete-variant), peaks with their sharpness blurred by both mapping (CN-induced) error and pointwise changes to gamete-variant sites.

Hence, to increase the signal to noise ratio, we subset only those scaffolds that showed no signs of CNV (**SM Figure 7**). With this filter only we plot the distribution of the minor allele frequencies of the two males. While analysis of BH3-2 showed a similar pattern as before, the signal of the Afus1 individual was much clarified (**SM Text 7**). Furthermore, this method allowed us to independently estimate the fraction of sperm in springtail bodies. The estimates are lower (30.39% and 33.96%) compared to the two-tissue model (35% and 38%). While we provide no explanation for the difference, we speculate the pileup method of estimating $f_h$ is noisier compared to the one based coverage, as the method of estimating 1n and 2n mapping coverages is extremely robust.

### SM Text 7: Noisier but supporting evidence in the reference male

Our study design contains only two male samples of the (potentially) PGE globular springtail *A. fusca* - Afus1, the deeply sequenced reference sample used to generate the assembly, and BH3-2, the resequencing sample. While BH3-2 shows very clear signals both in SNP (**Figure 4**) and pileup analyses (**SM Figure 8**), the reference male showed only extremely few heterozygous loci on autosomes we could use.

This could be caused by using the same reads for both assembly and SNP calling, as heterozygous loci can cause fragmentation of assemblies. Together with our threshold for assigning to chromosomes only scaffolds longer than 10,000 nt (**SM Text 1**) we could have filtered the majority of the scaffolds where Afus1 carried any heterozygous loci. However, heterozygosity of the species is not very extreme (**SM Figure 5**) to cause major problems during the assembly, therefore it is more likely the individual is simply very homozygous and the stringent SNP calling process did not pick enough true positive SNP calls to provide us a conclusive answer. If there is however a fraction of true positive SNP calls among the noisy SNP calls, they also show the same separation of distributions (**SM Figure 9**).

This idea is further supported by the pileup analysis (**SM Text 6**). Although we still observe a noisier signal compared to the BH3-2 individual, Afus1 individual also shows the signature of skewed coverage ratio between minor and major alleles of heterozygous loci on the autosomes, consistent with the PGE model (**SM Figure 8**).

Further investigation with a less fragmented reference genome would be needed to investigate further heterozygosity distribution across the genome in this sample.

### SM Figure 1: Mapping coverages of all resequencing samples

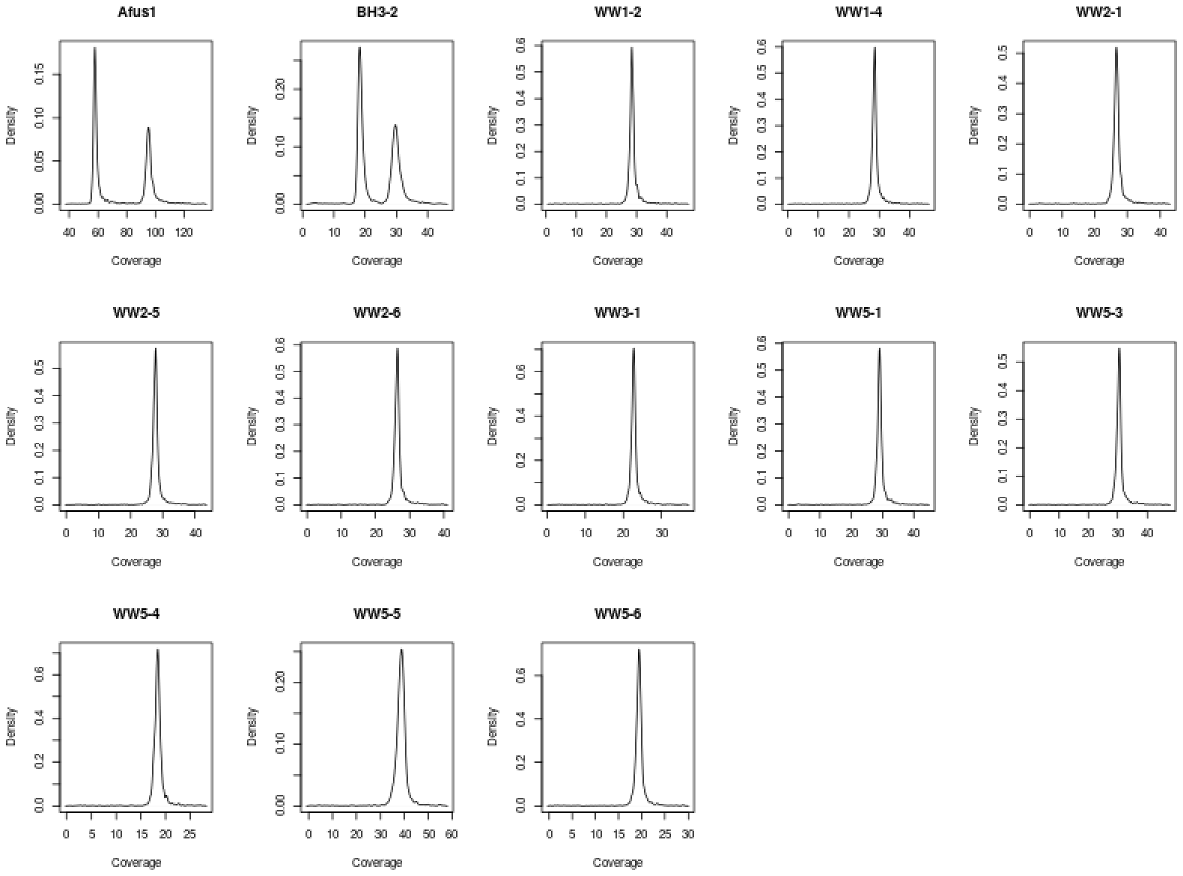

**SM Figure 1: Mapping coverage distributions of all sequenced individuals.** Mapping coverage distributions (see **Box 1**) estimated via kernel smoothing (see **Methods**). Coverage was used to determine the sex of the sample - male samples are expected to display two distinct peaks, females are expected to show just one. Besides the reference male (Afus1), only one more sample is male (BH3-2), all remaining sequenced samples were females.

### SM Figure 2: Sequencing coverages of males.

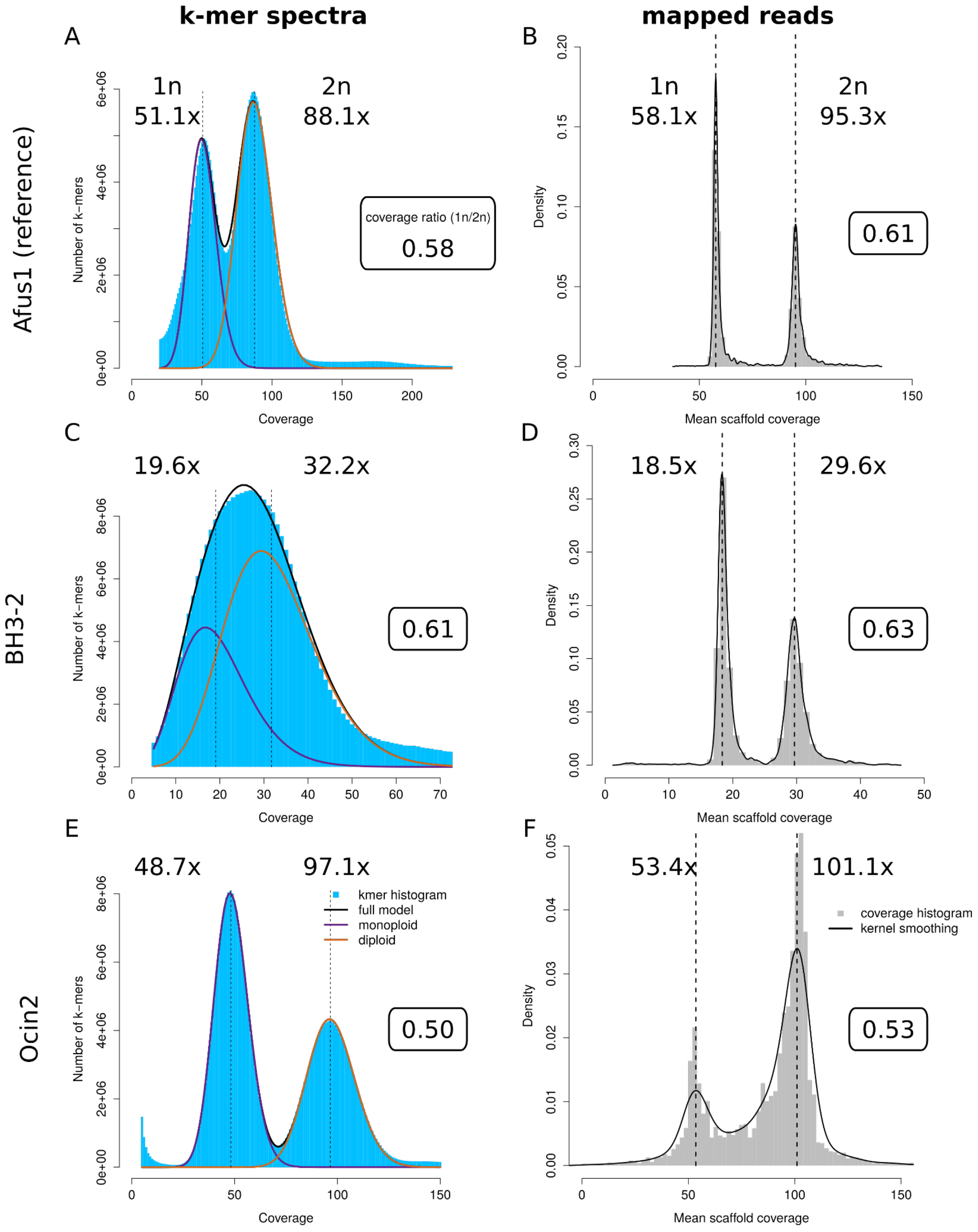

**SM Figure 2**: **Sequencing coverages of males.** The 1n peak represents X-linked regions, and in the case of k-mer spectra (A, C, E) also heterozygous autosomal positions. The second column of distributions show weighted scaffold coverage distributions from mapped reads. Panels A, B, C, D represent the reference (A, B) and resequenced (C, D) *A. fusca* males. In both cases 1n peak does not correspond to ½ of 2n peak. Compared to *O. cincta* male (E, F) with both kmer spectra and mapped coverages showing usually 1n and 2n peaks.

#### SM Figure 3: The effect of mixed tissue library on coverage

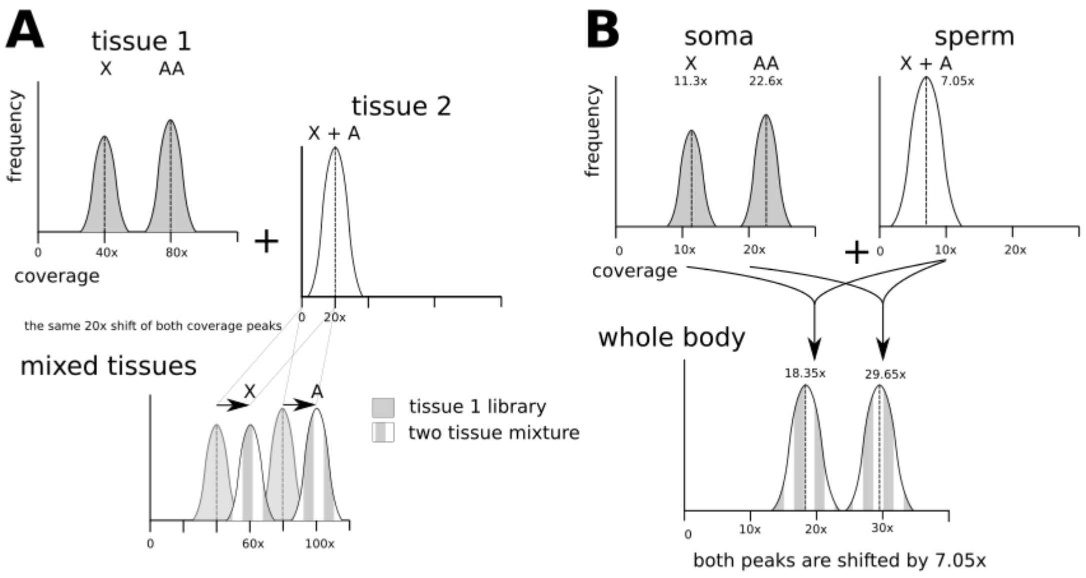

**SM Figure 3: The effect of mixed tissue library on coverage. A,** A hypothetical scenario of 2:1 mixture of tissue 1 and tissue 2 with two different karyotypes. The tissue 2 has the same ploidy for all the chromosomes (a single peak) and mixing with the tissue 1 will lead to a constant shift (symbolized by the arrow) that causes the peaks to be unevenly spaced in the mixed library (60x vs 100x). **B,** A mixture of the same two tissue types (AAX0 and AX karyotypes) with numbers that would explain the observed coverage of **BH3-2** (Figure 3)**.**

### SM Figure 4: Scheme of all considered two tissue models

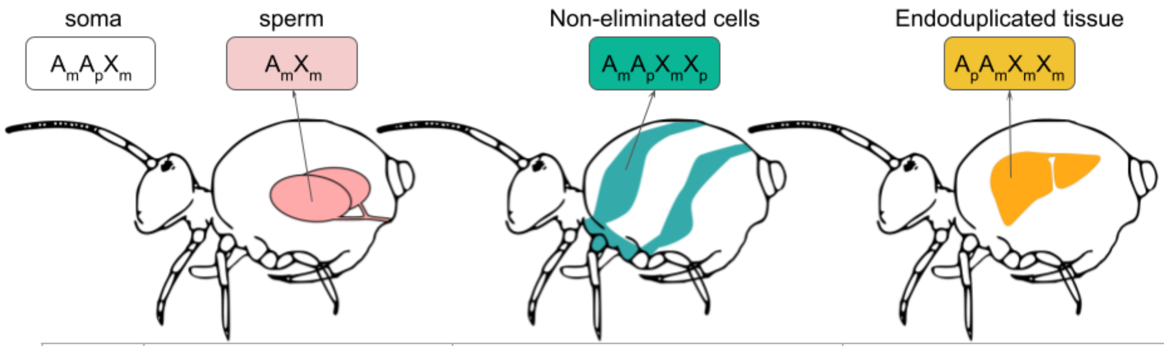

| | PGE | Imperfect elimination | endoduplication |
|---|---|---|---|
| $X_m$ | 17.44x | 14.365x | 17.44x |
| $X_p$ | - | 3.075x | - |
| $A_m$ | 17.44x | 14.365x | 14.365x |
| $A_p$ | 11.29x | 14.365x | 14.365x |

**SM Figure 4: Scheme of all considered two tissue models.** PGE is the model presented in the body of the manuscript. Non-eliminated cells and Endoduplicated tissue are models considered in **SM Text 4**. The coverage expectations are calculated for individual BH3-2.

### SM Figure 5: Female allele coverage supports.

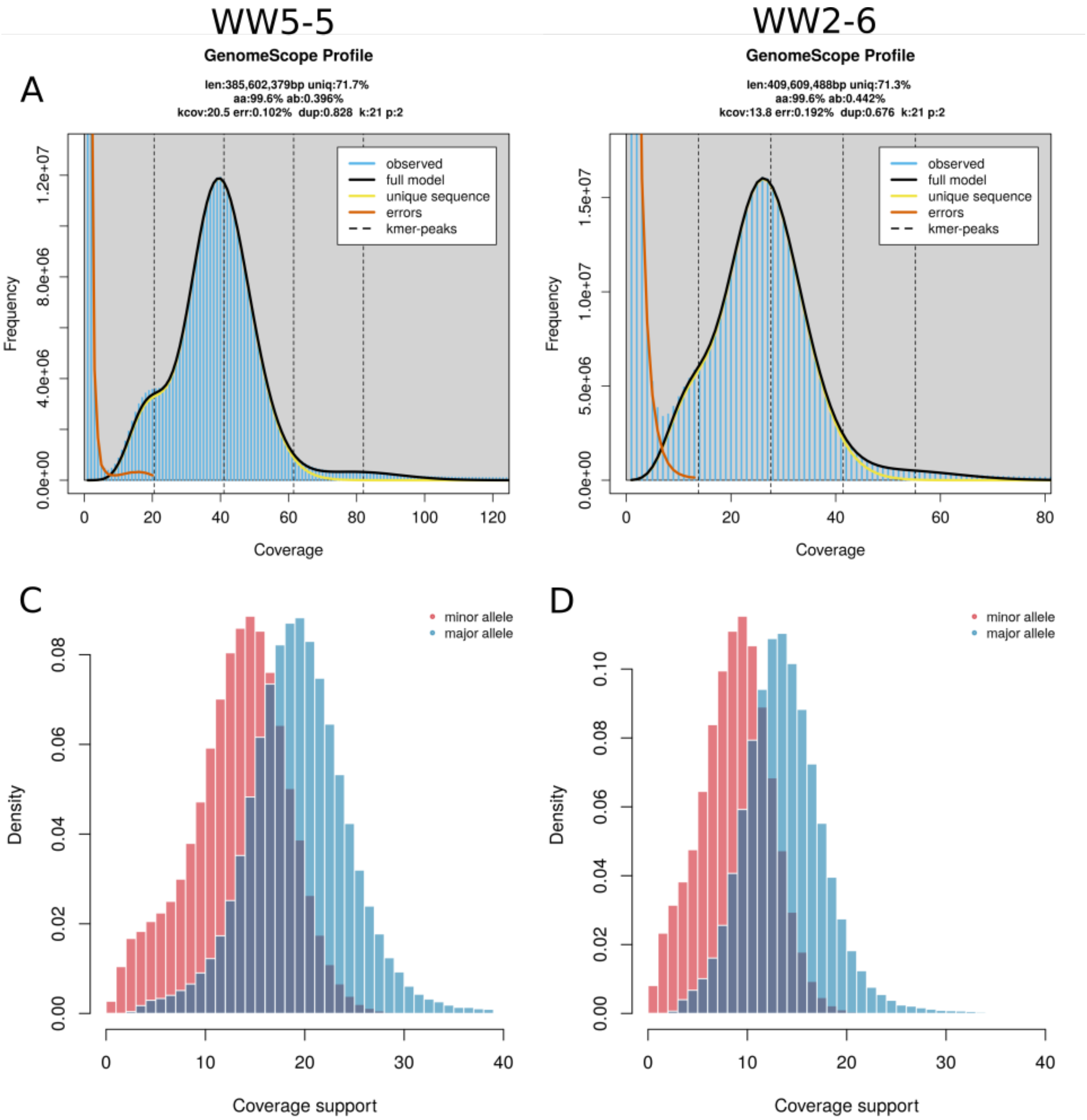

**SM Figure 5: Profiles and allelic supports of female *A. fusca.*** Available sequences of female Allacma fusca show now sign of 1n coverage peak shift. Here, the 1n peak represents solely heterozygous loci, as females carry all chromosomes in diploid state. However, autosomal heterozygous variants are still possible to decompose to “major” and “minor” alleles. The distributions are not very well separated as found in the male library (**Figure 4C**) and therefore the decomposition likely follows the same principle as in non-PGE species (shown in **Figure 4B**).

### SM Figure 6: Minor coverage ratios of heterozygous autosomal SNPs

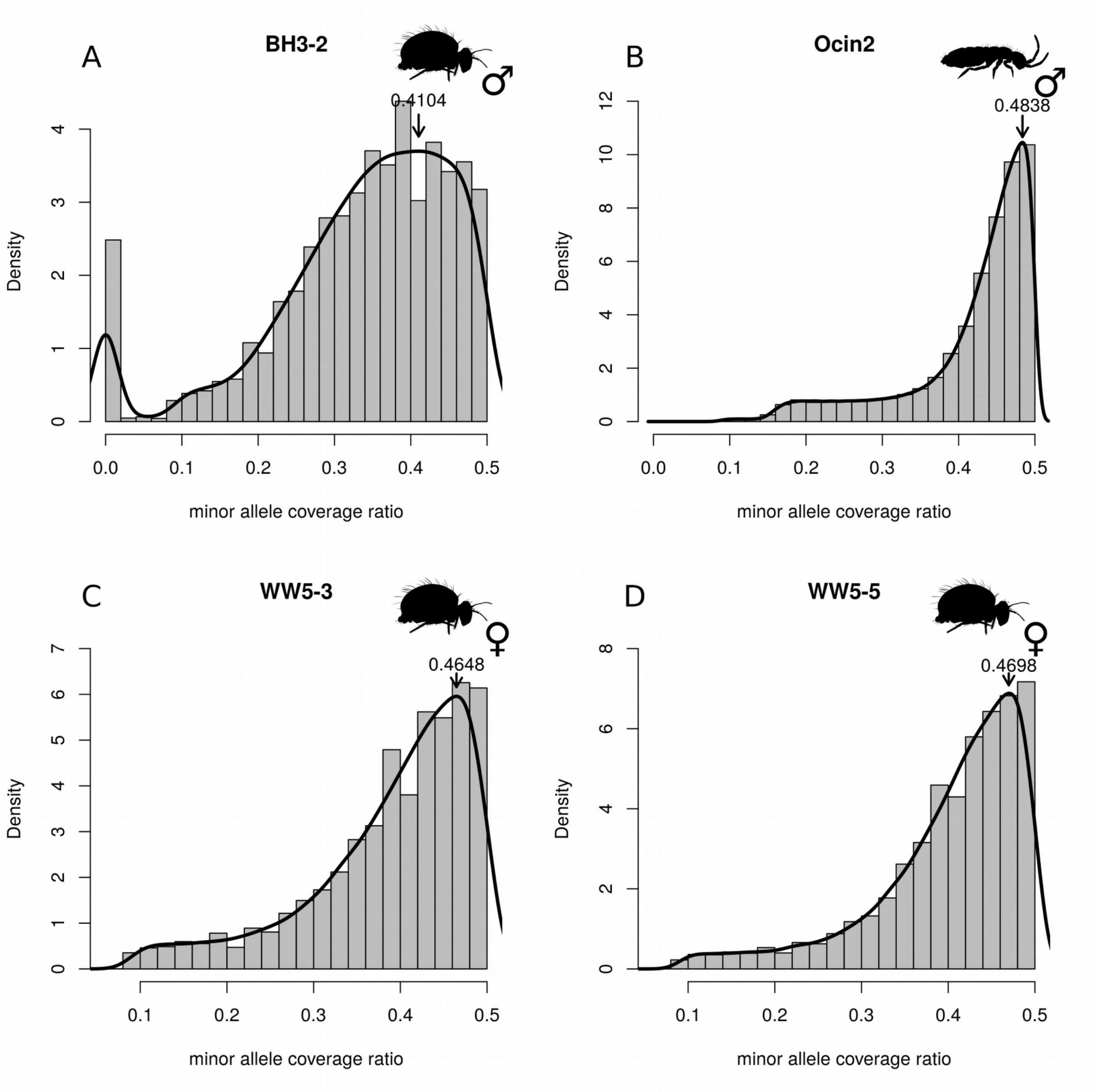

**SM Figure 6: Coverage ratios of autosomal heterozygous SNPs.** The resequenced individual (BH3-2, panel **A**) shows the peak of minor coverage distribution ($p_p$) at 0.4104, which corresponds to 30.39% of sperm in the springtail body and clearly shows a deviation from 0.5 coverage ratio expectation of maternal and paternal alleles. For balanced coverage ratio it is impossible to separate maternal and paternal alleles using short read data and therefore major and minor alleles will not really correspond to the parent of origin. The ratio of major and minor allele however still should be relatively close to 0.5, which was the case for both the non-PGE springtail (**B**) and two females analysed (**C** and **D**). Note female samples have a comparable coverage to BH3-2 (**SM Figure 1**). See **SM Figure 8** for comparison to the same plot created on raw pileup files.

### SM Figure 7: Subsetting scaffolds with low coverage variance

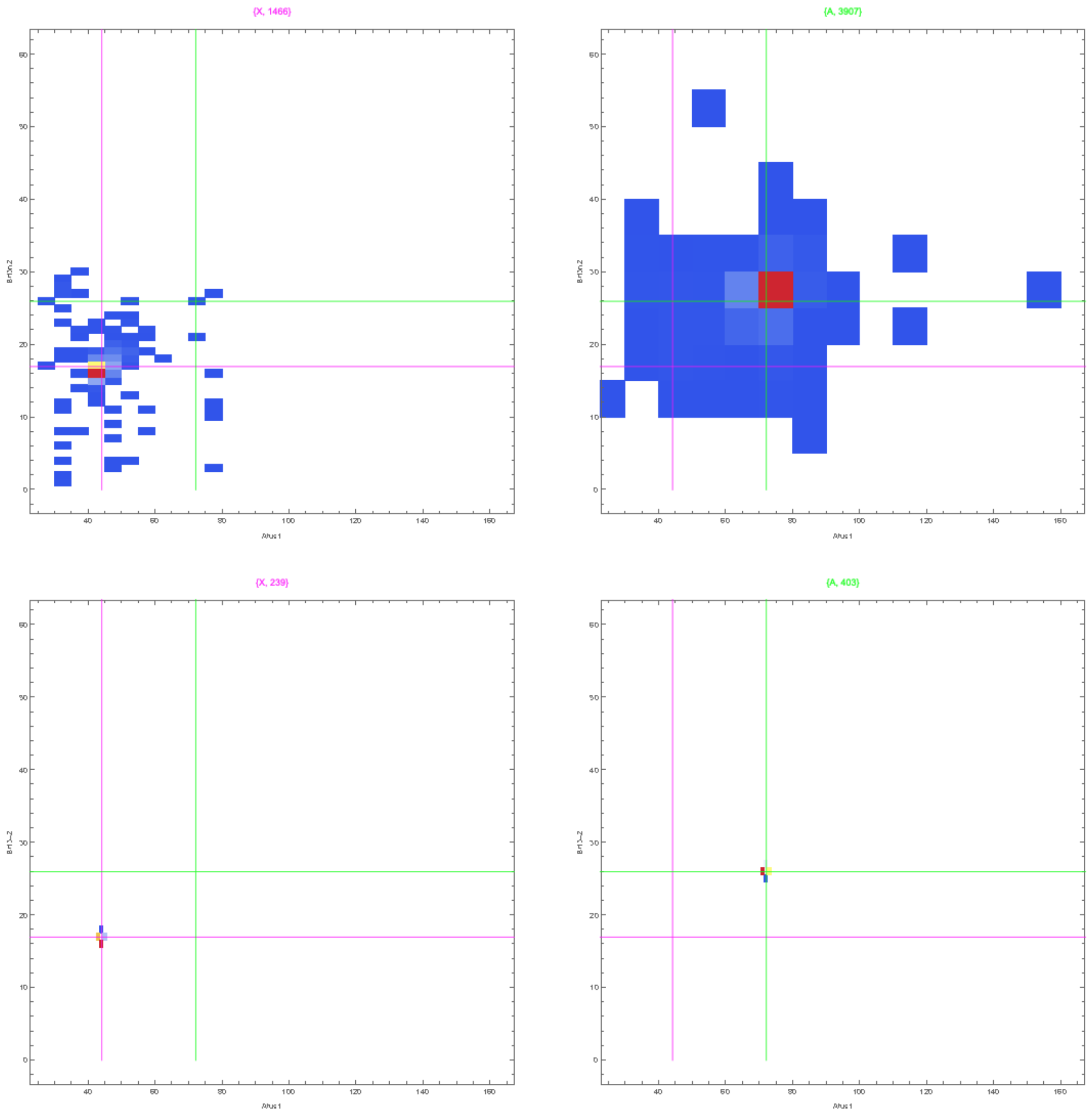

**SM Figure 7: Subsetting scaffolds to those with low coverage variation.** Top two panels are showing the coverage of the two male individuals for assigned X (left) and autosomal (right) scaffold. Bottom two panels are showing the subset scaffolds with no signs of copy number variation used for plotting of **SM Figure 8.**

### SM Figure 8: Estimated $f_h$ from raw pileups of reads

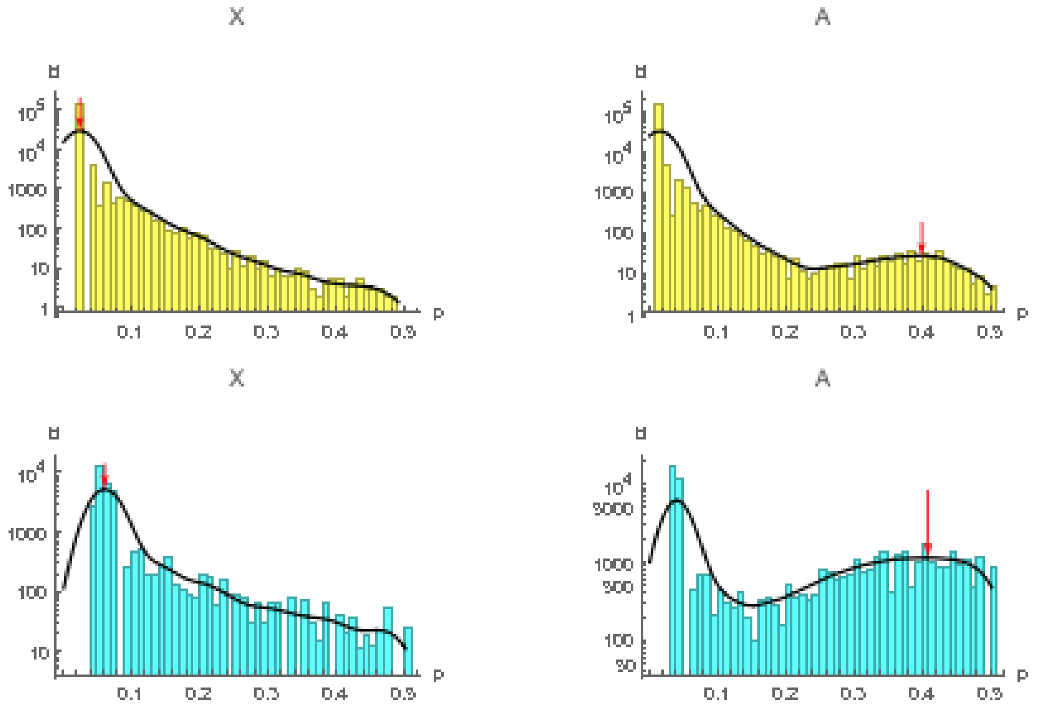

**SM Figure 8: Allele frequency in coverage pileups.** In this analysis we treat coverage ratios of all the bistates (genomic reference positions with two different nucleotides detected) as allele frequencies among the DNA molecules sequenced. As there was no SNP calling involved, sequencing errors will show up as bi states as well, therefore we search for additional peaks in the distribution. The X chromosomes showed no “peak” suggesting all the bistates represent sequencing or mapping errors. However, we observe a peak for coverage ratios 0.397 in Afus1 (yellow) and 0.406 in BH3-2 (teal). Note even the reference male with low heterozygosity levels shows some detectable coverage ratio peak, although the signal is much weaker compared to BH3-2. The estimated paternal allele frequencies $p_p$ correspond to 33.96% of the fraction of sperm in the body ($f_h$) for Afus1 and 31.39% in BH3-2 respectively. See **SM Figure 6A** for comparison to a plot created on using coverage support of called SNPs.

SM Figure 9: Allele coverage supports in the reference male (Afus1).

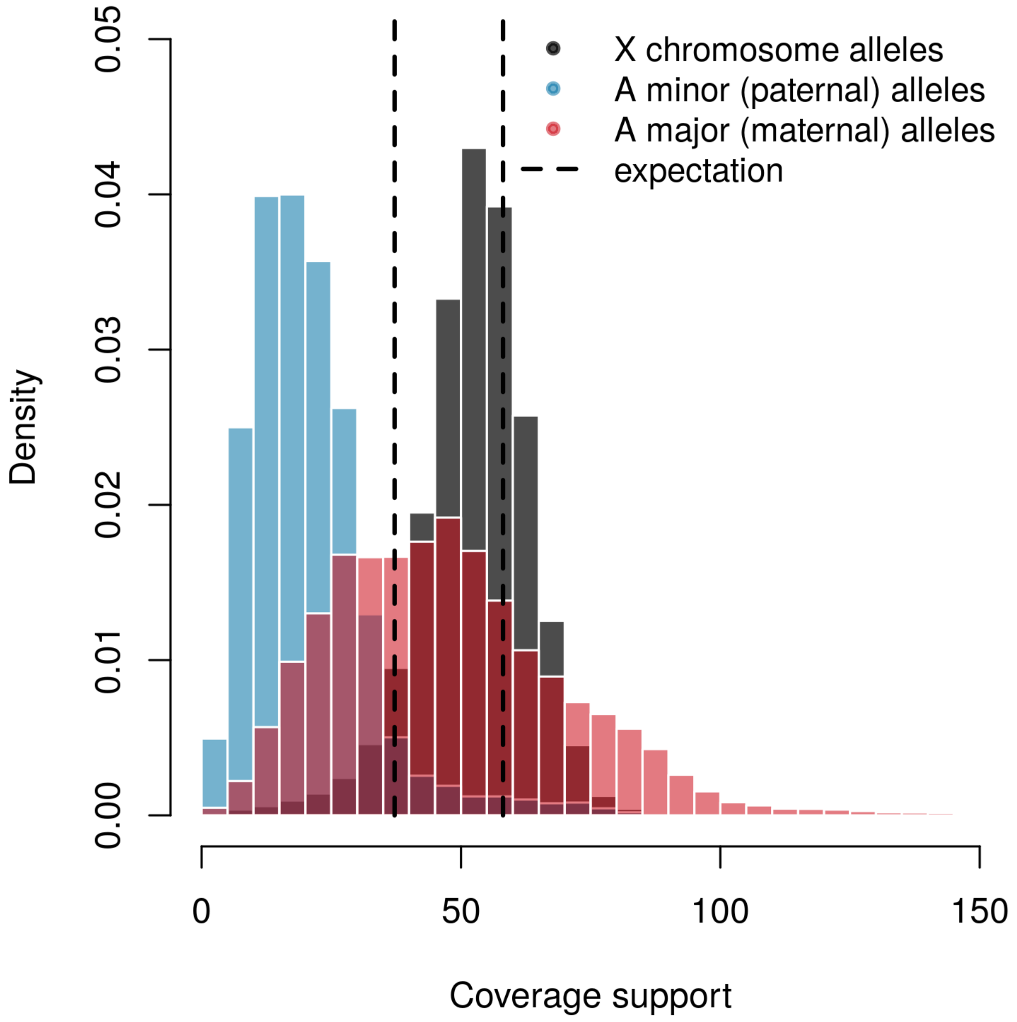

**SM Figure 9: Allele coverage supports in the reference male (Afus1).** Reference male should in theory show the same pattern as the individual BH3-2 (Figure 3C). However, there were a lot fewer called heterozygous variants and majority of the minor allele coverages showed a lot lower coverage than expected (in dashed lines). The major allele frequencies show extremely high coverage variance, riding additional red flag regarding the detected variants. The X chromosome alleles are homozygous reference alleles located at all positions on X chromosomes where at least one was variant detected in any of the resequencing individuals.

#### SM Figure 10: Power analysis

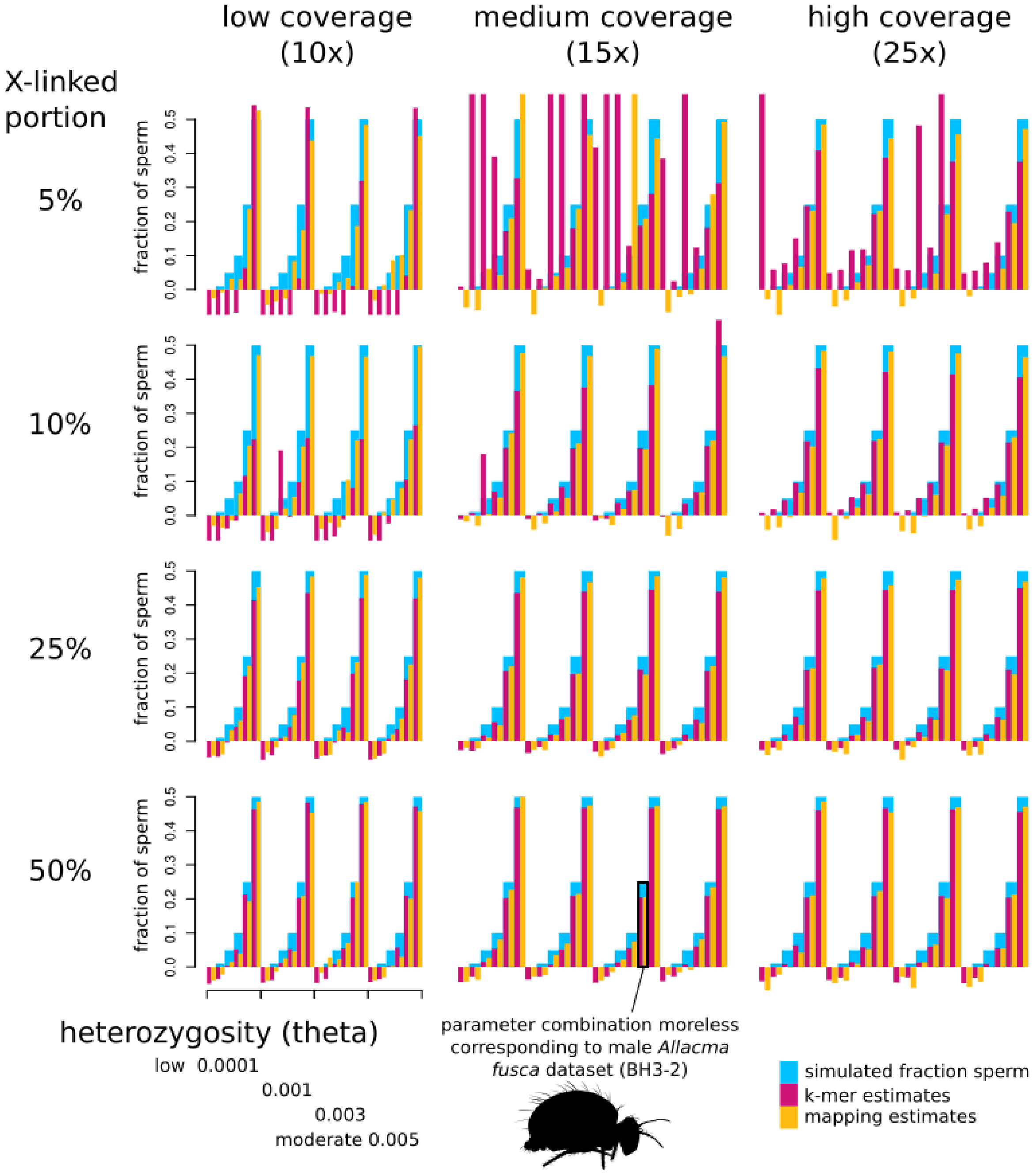

**SM Figure 10: Power analysis of the two tissue model.** We simulated 20Mbp long genomes with X-linked portions spanning 5 - 50% of the total length (rows of panels). We simulated three coverage levels (per maternal haplotype, three columns panels) and four levels of heterozygosity (expressed as theta, 4 subdivisions of each panel). For each combination of parameters we simulated 6 levels of sperm in the sample assuming PGE model (y-axis, blue bars). The colored bars are the inferred fractions of sperm using kmers (in red) and via mapping (in yellow). The combination of parameters corresponding to BH3-2 is highlighted with a black rectangle. Estimates that are very far from expectations are due to poor convergence of the two tissue models. Note overall the two tissue model underestimates the simulated fraction of sperm.

#### SM Table 1: Literature with evidence for PGE.

https://docs.google.com/spreadsheets/d/1Is_KKSrFNzEzdT-o0Pdj-z4Hh2uC82XXFPzP21SeFXI/edit?usp=sharing

#### SM Table 2: Table of variants called in all male samples.

The grey lines indicate assembly spans and the span of the scaffolds with autosomal and X-linked chromosomal assignments (rounded in Mbp). While lines give the post-filtering number of variants found in whole assemblies, autosomes and X chromosomes in heterozygous (0/1) or homozygous (1/1) state. Heterozygous (0/1) variant calls on X-linked scaffolds represent false positives as only males are included in the table. Most of the heterozygous alleles in the reference individual did not anchor to chromosomes (see **SM Text 7** for more details).

| species | | Total assembly span | | Autosomal scfs | | X-linked scfs | |
|---|---|---|---|---|---|---|---|
| | sample | 0/1 | 1/1 | 0/1 | 1/1 | 0/1 | 1/1 |
| *Allacma fusca* | | 426 Mbp | | 93 Mbp | | 78 Mbp | |
| | Afus1 (ref) | 660,165 | 6,332 | 21,471 | 242 | 6,599 | 194 |
| | BH3-2 | 761,870 | 457,721 | 227,570 | 117,693 | 7,731 | 60,999 |
| *Orchesella cincta* | | 286 Mbp | | 224 Mbp | | 63 Mbp | |
| | Ocin2 | 3,190,875 | 1,910,065 | 1,959,258 | 915,879 | 144,204 | 400,001 |

### SM Table 3: Table of fractions of sperm estimated by various techniques.

The table of estimated fractions of sperm for five *A. fusca* samples (PGE springtail, 2 male and 3 female) and *Orchesella cincta* male sample. Only the two *A. fusca* males are expected to have any significant portion of body to be sperm with a different karyotype. While the reference free technique (k-mer spectrum) estimated low fractions for 3 out of 4 control samples, the mapping coverage resulted in ~10% estimated fraction of sperm in Ocin2 sample. This was caused most likely by assembly errors that merged together X-linked and autosomal contigs which artificially caused the two coverage peaks (1n and 2n) to be closer to each other than expected. The single female sample with high negative proportion of estimated fraction of sperm is due to extremely low coverage (see power **SM Text 5** analysis for more details). SNP calling seems to estimate a very high fraction of sperm in Afus1, that is likely due to a very high fraction of false positives among heterozygous autosomal SNPs. With exception of this one estimate, all remaining estimates of the fraction of sperm in ***A. fusca*** males are relatively consistent and range between 27.5 - 38.4%.

| species | sample | sex | Expected shift | K-mer spectrum | Mapping coverage | SNP calling | Analysis of pileups |
|---|---|---|---|---|---|---|---|
| *Allacma fusca* | Afus1 | ♂ | Yes | 27.5% | 35.0% | 57.1% | 34% |
| | BH3-2 | ♂ | Yes | 35.3% | 38.4% | 30.4% | 31.4% |
| | WW5-3 | ♀ | No | -6.6% | - | 13.2% | - |
| | WW5-5 | ♀ | No | -6.8% | - | 11.4% | - |
| | WW2-6 | ♀ | No | -23.2% | - | 13.2% | - |
| *Orchesella cincta* | Ocin2 | ♂ | No | 0.6% | 10.3% | 6.3% | - |
